# Antiviral RNA interference targets viral transcripts but not genomes of RNA viruses in *Drosophila melanogaster*

**DOI:** 10.1101/2024.04.10.588985

**Authors:** Emanuele G. Silva, Álvaro G. A. Ferreira, Thiago Henrique L. Jiran, Isaque João S. de Faria, Juliana N. Armache, Siad C. G. Amadou, Gabrielle Haas, Franck Martin, Jean Luc-Imler, João T. Marques

## Abstract

RNA interference (RNAi) mediated by the small interfering RNA (siRNA) pathway is a major antiviral mechanism in insects. This pathway is triggered when double-stranded RNA (dsRNA) produced during virus replication is recognized by Dicer-2, leading to the formation of virus-derived siRNA duplexes. These siRNAs are loaded onto the programmable nuclease Argonaute-2 (AGO2), with one strand serving as a guide to target and cleave fully complementary sequences of viral RNAs. While siRNAs are generated from viral dsRNA, the specific viral RNA species targeted for silencing during RNA virus replication remains unclear. In this study, we characterized the primary viral RNA targets of the *Drosophila* siRNA pathway during infections caused by negative and positive RNA viruses, namely Vesicular stomatitis virus (VSV) and Sindbis virus (SINV). Our findings reveal that polyadenylated transcripts of VSV and SINV are the major targets of silencing by the siRNA pathway during infection, likely when they are poised for translation. Consistent with earlier findings, we confirmed that AGO2 copurifies with ribosomes, and this is not affected by virus infection. Therefore, we propose that the inhibition of the replication of RNA viruses in *Drosophila* results from the silencing of incoming viral transcripts, facilitated by the association of AGO2 with ribosomes.

**Author Summary:** The small interfering RNA (siRNA) pathway mediates major antiviral immune response in insects, functioning to cleave viral RNA. While this pathway has been extensively studied in the fruit fly *Drosophila melanogaster*, the specific molecular targets of inhibition by the siRNA pathway have remained unclear. In this study, we aimed to elucidate these targets. Our findings demonstrate that polyadenylated transcripts produced during viral infection are the primary targets of the Drosophila siRNA pathway, in the case of both negative and positive single-stranded RNA viruses. The silencing of these transcripts accounts for the antiviral effect of the siRNA pathway, suggesting that direct targeting of viral RNA genomes is unlikely to occur. We confirmed that Argonaute-2 (AGO2), the core component of the silencing complex, co-purifies with ribosomes and also show that this association is not affected by viral infection. This suggests that AGO2 is in permanent association with ribosomes where it can efficiently scan viral transcripts before they undergo translation by the cellular machinery, thereby preventing viral replication. These results provide valuable insights into the mechanism of gene silencing the siRNA pathway.

## Introduction

The small interfering RNA (siRNA) pathway mediates a major antiviral defense in *Drosophila* and other insects (1). This RNA interference (RNAi) pathway is triggered when the RNA helicase and RNase III enzyme Dicer-2 (Dcr-2) senses dsRNA generated during viral infection. Upon recognition, Dcr-2 processes dsRNA into siRNAs that are loaded onto Argonaute-2 (AGO2), a programmable nuclease, to form the RNA-induced silencing complex (RISC), which uses the sequence of the siRNA to find complementary targets (1,2). This highly specialized antiviral siRNA pathway was extensively characterized in the fruit fly *Drosophila melanogaster* (3–6). Flies deficient for core proteins of the siRNA pathway, namely AGO2, Dcr-2 and R2D2, are more susceptible to a range of viruses with RNA or DNA genomes, such as Flock house virus (FHV), Sindbis virus (SINV), Vesicular Stomatitis viruses (VSV) and Invertebrate iridescent virus 6 (IIV6) (5–9).

Flies infected with positive and negative stranded RNAs viruses accumulate virus-derived siRNAs (vsiRNAs) along the whole genome, on both strands, indicating they are derived from dsRNAs intermediates of replication (7). Thus, these vsiRNAs have the potential to target any viral RNA species in infected cells, both genomes and transcripts. Notably, silencing of either genomes or transcripts will affect virus replication and indirectly inhibit the production of all types of viral RNAs, which makes it hard to determine the initial target. In the case of *Tospovirus* infected plants, where RNAi is also a potent antiviral defense, viral transcripts were shown to be the primary target of silencing, in agreement with the fact that viral genomes are protected by ribonucleoproteins (10). However, since the mechanism of antiviral RNA interference is considerably different between plants and animals, it is unclear how generalizable this observation is. Our previous work has suggested that the polyadenylated transcripts are preferentially targeted by RISC in infected *Drosophila* but this remains to be formally demonstrated (7).

Here, we employed the well-characterized insect model, *Drosophila melanogaster*, to identify the preferential targets of the siRNA pathway during infection with negative- and positive-stranded RNA viruses, VSV and SINV, respectively. Our findings suggest that antiviral RNA interference in flies operates by targeting viral transcripts precisely when they are poised for translation by cellular ribosomes.

## Results

### Intermediates of replication are required to induce an antiviral response against VSV

VSV is a negative-stranded RNA virus of the *Rhabdoviridae* family that induces a potent siRNA response in *Drosophila* (7,9). In order to evaluate the specificity and persistence of antiviral responses in *Drosophila*, we developed an *in vivo* strategy where flies were primed with replication competent and UV-inactivated VSV before being exposed to a secondary infection with the same virus. The general strategy is presented in **Fig 1A**. To differentiate primary and secondary infections, we used a recombinant virus expressing GFP (VSV-GFP) on the latter. We then monitored expression of GFP as a reporter for the second virus compared to the VSV L gene encoding the viral polymerase that reflects replication of both primary and secondary infections. We observed no significant changes in the replication of VSV-GFP in flies primed with UV-inactivated VSV but GFP expression given was strongly inhibited by prior exposure to the replication competent virus **(Fig 1B)**. Although there are many possibilities to explain this difference, we reasoned that dsRNA intermediates of genome replication would only be generated by replication competent viruses. The dsRNA generated by the first infection would then be able to trigger the siRNA pathway and restrict replication of the second homologous virus. To address this hypothesis, we synthesized dsRNA targeting the nucleoprotein gene (N) from VSV (dsVSV-N) **(Fig 1C and S1)**. We also synthetized dsRNA targeting an unrelated gene from *Firefly luciferase* as a control of dsRNA treatment (dsFluc) (Fig 1C and S1). dsRNAs were made *in vitro* from cDNA templates targeting regions of virus genes and the control Fluc and were tracked *in vivo* for their efficiency.

**Fig 1.**
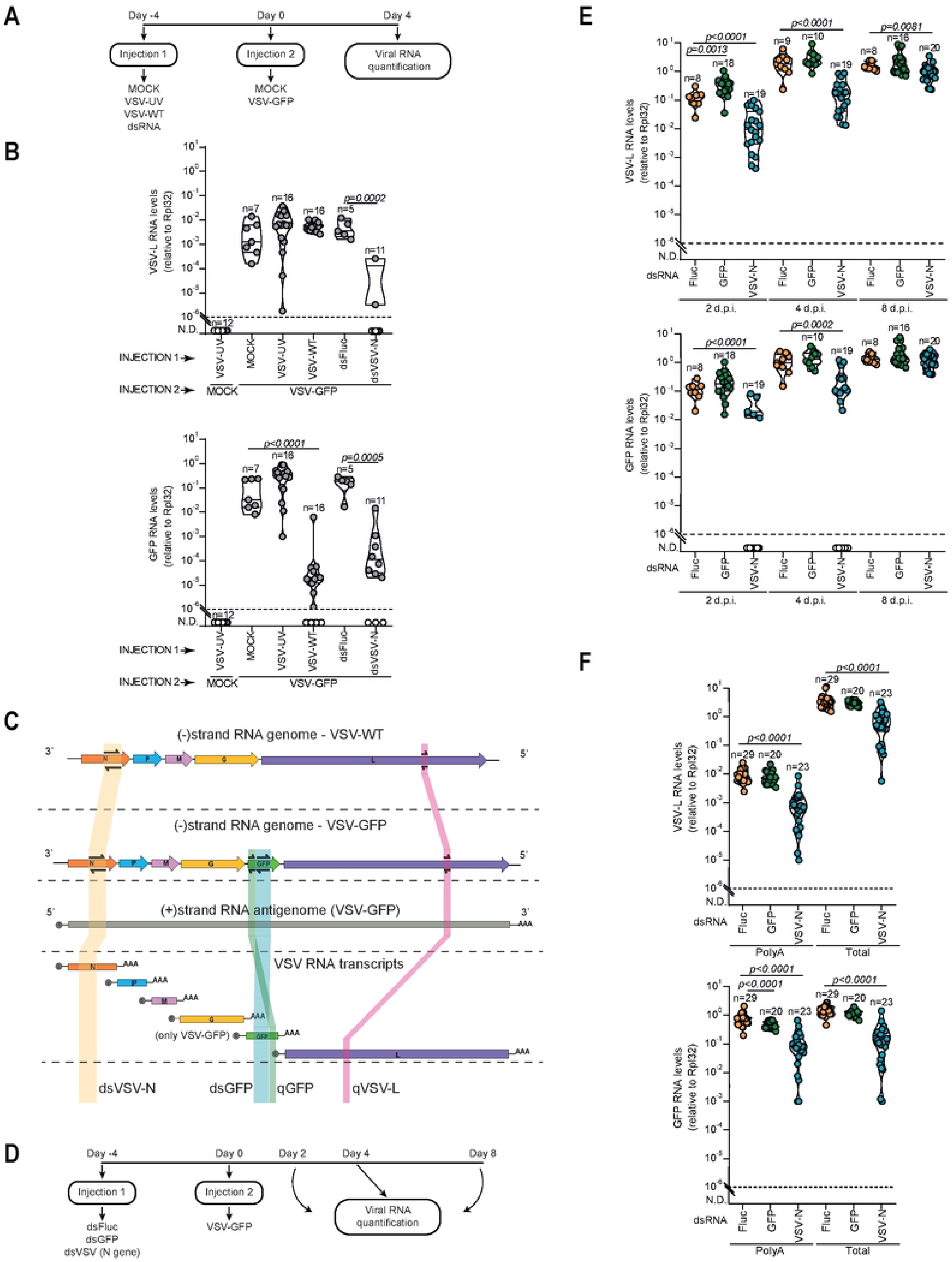
Polyadenylated viral transcripts are preferential targets for dsRNA mediated silencing during VSV infection. **(A**,**D)** Experimental design of dsRNAs treatment and VSV infections in *Drosophila*. **(B)** VSV-L and GFP RNA levels at 4 days post infection with VSV-GFP. **(C)** Overview of VSV-WT and VSV-GFP RNA species and the target regions selected for primers design used for dsRNA synthesis (dsVSV-N and dsGFP) and RT-qPCRs assays (qGFP and qVSV-L). **(E)** VSV-L and GFP RNA levels at the indicated times post infection with VSV-GFP. (**F)** Levels of polyadenylated and total RNAs corresponding to VSV-L and GFP genes at day 4 post infection with VSV-GFP. RNA levels were assessed by RT-qPCR performed in triplicate and normalized to Rpl32 mRNA levels. Violin plots show the frequency of data distribution with the median and quartiles. Each dot represents an individual sample. Numbers of samples are indicated above each violin plot. The dashed line represents the detection threshold of RT-qPCR. *p* values are shown for statistical significant *(p<0*.*005)* determined by two-tailed Mann–Whitney U-test.

Next, we injected these dsRNAs into *Drosophila* adults and infected them with VSV-GFP 4 days later. Treatment with dsVSV-N induced ∼ 1000-fold decrease in the levels of the VSV L gene and GFP RNAs compared to dsFluc-treated flies **(Fig 1B)**. Taken together, our results suggest that the protection of flies triggered by replication competent VSV is likely mediated by the detection of dsRNAs by Dcr-2, which result in the production of virus-specific siRNAs that control infection by homologous viruses.

### Polyadenylated RNAs generated during VSV infection are preferential targets of siRNA pathway

Since synthetic dsRNA targeting VSV primes antiviral RNAi similar to a primary infection, we decided to use our model to further understand how the siRNA pathway targets viral replication. Fig 1C shows a schematic representation of the different RNA species produced during VSV infection. The viral RNA dependent RNA polymerase (RdRp) encoded by the L gene binds the encapsidated genome at the 3’ leader region, producing capped and polyadenylated transcripts for each viral gene in the order that they appear from 3’ to 5’, Nucleoprotein (N), Phosphoprotein (P), Matrix (M) protein, Glycoprotein (G), and Large polymerase (L) (11–13). The RdRp also transcribes the antigenome of opposite polarity, which is a template for new the synthesis of new viral genomes. Genome replication starts when enough nucleoprotein is present to encapsidate neo-synthetized genomes (12).

In our model, GFP reflects viral replication, but production of the protein is not essential for replication. Thus, if GFP transcripts are targeted by the silencing machinery, this would not affect VSV replication. In contrast, if viral genomes or antigenomes were to be targeted, silencing of the GFP within the viral genomic RNA would block replication. This allowed us to test which species of viral RNAs were targeted during antiviral RNA interference in *Drosophila*. Thus, we injected dsRNA targeting GFP (dsGFP), dsVSV-N and dsFluc into distinct groups of flies before VSV infection and collected them at different time points to access viral loads **(Fig 1D)**. As observed before, dsVSV-N strongly inhibited VSV replication, as shown by decreased expression of the VSV-L gene and GFP throughout the kinetics of infection **(Fig 1E)**. In contrast, treatment with dsGFP did not decrease levels of VSV L **(Fig 1E)**. These results suggest that siRNAs generated from dsRNA processing did not target the VSV-GFP genome since it did not decrease virus replication. Notably, dsGFP did not significantly decrease levels of the GFP RNA in VSV-GFP infected cells **(Fig. 1E)**, which we attribute to the fact that GFP levels are a combination of transcripts and RNA genomes. Thus, we next measured the abundance of polyadenylated VSV RNAs to directly evaluate levels viral transcripts. Similar to the results measuring total RNA, levels of polyadenylated RNAs corresponding to L gene and GFP were significantly reduced by treatment with dsVSV-N **(Fig 1F)**. In contrast, dsGFP treatment did not affect levels of VSV-L but reduced transcript levels of GFP, even though total levels were not significantly affected **(Fig 1F)**. These results reinforce that total GFP levels reflect the sum of transcript and genomic RNA, where the latter is more abundant. Taken together, these results indicate that viral transcripts and not genomic RNAs are the major target of siRNA pathway during VSV infection in *Drosophila*.

### Transcripts from SINV, a positive stranded RNA virus, are preferentially targeted by the siRNA pathway

To investigate whether our observations could be applied to other viruses, we used a positive stranded RNA virus, SINV, that belongs to the *Togaviridae* family. The single-stranded RNA genome of SINV also serves as an mRNA for the translation of nonstructural proteins (nsPs) (14). nsPs, including the RdRp, promote the synthesis of new genomes, antigenomes and a subgenomic RNA encoding structural proteins (sP) (15). In our experiments, we used a version of SINV that expresses GFP (SINV-GFP) as an independent subgenomic RNA that is dispensable for viral replication, similar to our recombinant VSV-GFP.

We analyzed the kinetics of SINV-GFP replication in flies injected with dsRNA targeting nsP4, encoding the RdRp of SINV (dsSINV-nsP4) and compared to injection of dsGFP and the unrelated dsFluc as a control **(Fig 2A,B and S1)**. We observed a 10 to 100-fold reduction in the viral load of SINV as reported by the levels of another SINV gene, nsP2 at all times post-infection. We also observed a reduction of the levels of SINV sP and GFP in flies treated with dsSINV-nsP4 after 2 days post-infection, indicative of general inhibition of viral replication **(Fig 2C,D)**. In contrast, dsGFP did not consistently affect the viral load of SINV as indicated by the levels of nsP2 or sP, but it significantly decreased the GFP subgenomic RNA at all time points **(Fig 2C)**. We observed a significant decrease in nsP2 levels in flies treated with dsGFP at 8 days post infection but it was not as robust as the effect of dsSINV-nsP4 **(Fig 2C)**. Thus, targeting GFP mostly affected its own expression levels but not SINV-GFP replication. These results reinforce the idea that viral transcripts but not genomes are the major targets for RNA silencing. Considering that the positive strand of the SINV genome can be directly translated as a transcript, it was unexpected to have such a small effect on viral replication when GFP was targeted. This indicates that dsGFP only triggered silencing of its own subgenomic RNA, leaving unaffected the full-length genomic RNA, suggesting that the silencing machinery may only be able to scan targets close to the ORF that is being translated.

**Fig 2.**
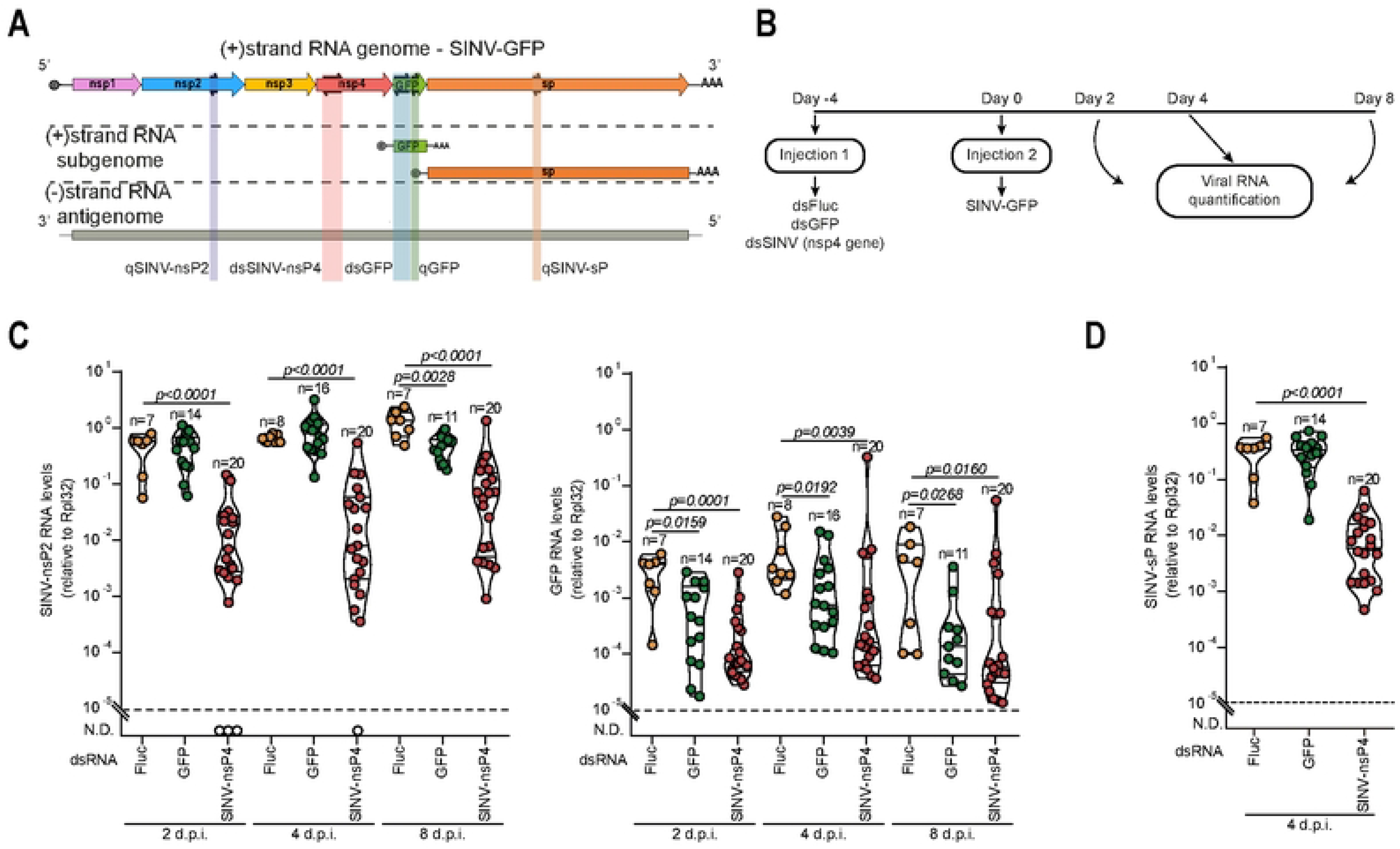
Translated RNAs are preferential targets for dsRNA mediated silencing during SINV infection. **(A)** Overview of SINV-GFP RNA species and the target regions selected for primers design used for dsRNA synthesis (dsSINV-nsp4 and dsGFP) and RT-qPCRs assays (qSINV-nsp2, qSINV-sp and qGFP). **(B)** Experimental design of dsRNAs treatment and SINV-GFP infections in *Drosophila*. **(C)** SINV-nsp2 and GFP RNA levels at the indicated times post infection with SINV-GFP. **(D)** SINV-sp RNA levels at day 2 post infection with SINV-GFP. RNA levels were assessed by RT-qPCR performed in triplicate and normalized to Rpl32 mRNA levels. Violin plots show the frequency of data distribution with the median and quartiles. Each dot represents an individual sample. Numbers of samples are indicated above each violin plot. The dashed line represents the detection threshold of RT-qPCR. *p* values are shown for statistical significant *(p<0*.*005)* determined by two-tailed Mann–Whitney U-test.

### AGO2 associates with ribosomes regardless of the infection

Previous studies in *Drosophila* S2 cells found that AGO2, the main component of RISC, is associated with cellular ribosomes (16,17). There, AGO2 would be able to scan viral mRNAs before they can be translated by the cell machinery. To confirm these observations during viral infection, we separated ribosome enriched (P100) and depleted (S100) fractions of S2 cells as described (18) **(Fig 3A)**. We used both control and VSV infected cells (lines 5 and 6) to analyze whether the virus could induce changes in AGO2 localization **(Fig 3B)**. In both infected and control cells, the ribosomal protein S15 (Rps15) was enriched in the P100 fraction, confirming that ribosomes were efficiently separated by our protocol **(Fig 3B)**. Regardless of the infection, AGO2 was present in the P100 sample (lines 2 and 4) and depleted from S100 fractions (lines 1 and 3), confirming that it is associated to ribosomes **(Fig 3B)**. Notably, we did not observe a significant difference in AGO2 accumulation in the ribosomal fraction during infection, suggesting that its localization is not regulated in response to the virus **(Fig 3B)**. Interestingly, in VSV infected cells, the viral RdRp was found both in ribosomal enriched and depleted fractions, possibly indicating multiple complexes responsible to the synthesis of the different viral RNA species **(Fig 3B)**. Altogether, our results suggest that AGO2 is closely associated with ribosomes where it can easily monitor incoming viral mRNAs to promote their degradation. This determines the preference of transcript silencing by the siRNA pathway during viral infection.

**Fig 3.**
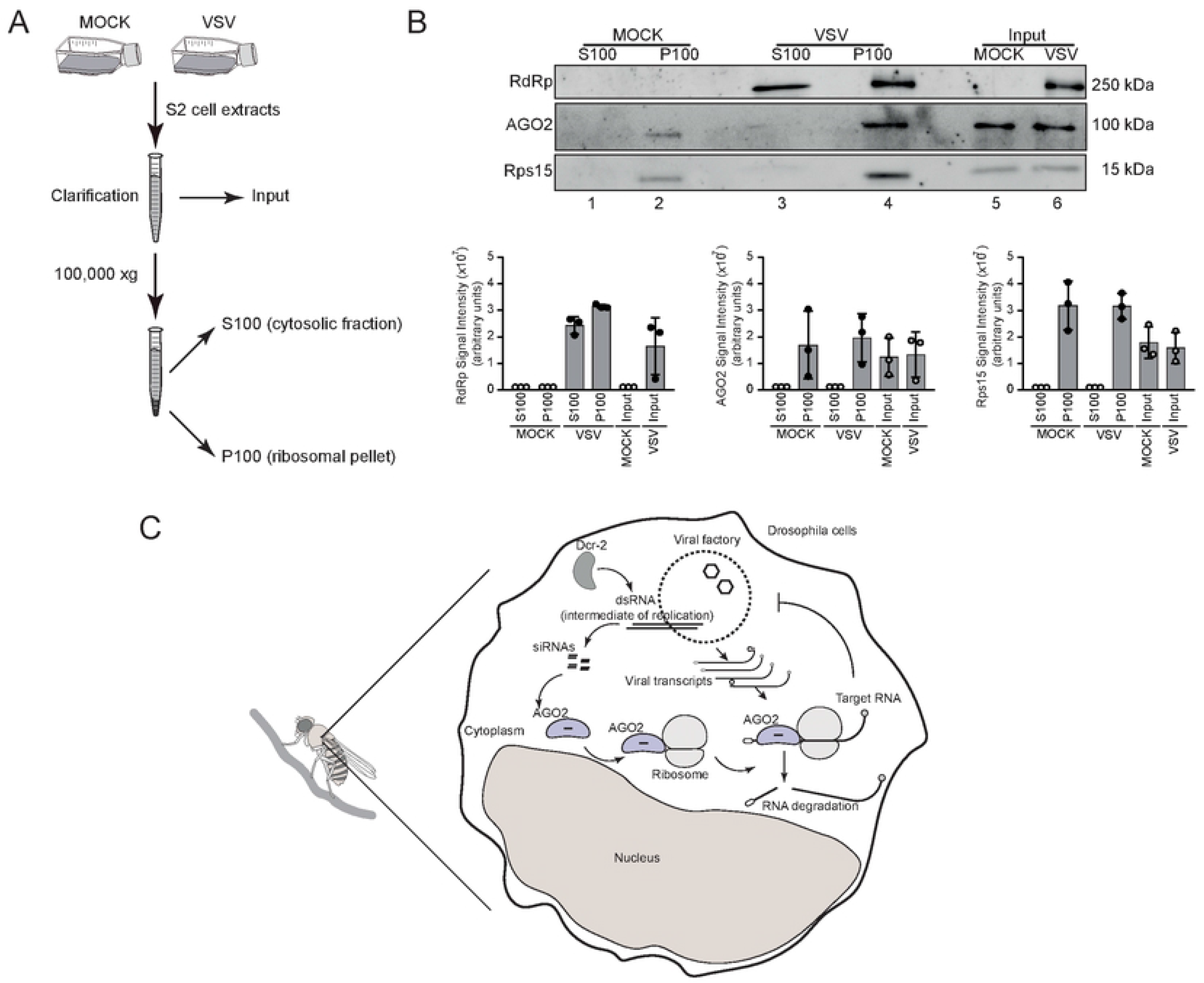
AGO2 association with ribosomes is not affected by virus infection. **(A)** Overview of the cell fractionation strategy. Mock- or VSV-infected cells lysate were submitted to high-speed centrifugation (100,000g). The S100 and P100 containing the cytoplasmic and ribosomes fractions, respectively, were extracted with high salt buffer and proteins were analyzed by Western blotting. (**B)** Representative Western blotting (upper panel) and quantification of signal intensity (lower panel) for VSV-RdRp, AGO2 and Rps15 proteins of S100 and P100 fractions and input. Graphs represent the means ± SD of three independents experiments indicated by the dots. **(C)** Overview of the molecular mechanism of inhibition by the siRNA pathway, which preferably targets RNA transcripts during infection with RNA viruses in *Drosophila*.

## Discussion

Although the antiviral siRNA pathway has been extensively studied in *Drosophila*, it remains unclear how it targets viral replication. Here, we have shown that viral RNA transcripts and not the RNA genome or antigenome are the major targets of the siRNA pathway during infection of RNA viruses in *Drosophila*. We propose that AGO2 and mature RISC are found in association with ribosomal proteins, where they can scan viral transcripts before they can hijack the cellular machinery **(Fig 3C)**. This is consistent with results from our previous study, where we showed that polyadenylated viral RNA from VSV and SINV were more strongly silenced in a Dcr-2 dependent manner compared to total RNAs (7).

Recently, Hong and collaborators showed that RISC-associated nuclease activity acts at the level of viral transcripts but not genomic RNA from Tospovirus in infected plants (10). They suggest that viral transcripts are preferentially targeted by the RNAi machinery. This could reflect the fact that viral transcripts are not protected in ribonucleoprotein complexes unlike viral genomes/antigenomes. Recent work has suggested that the limiting step in human AGO2 mediated mRNA degradation consist of unmasking target sites by translating ribosomes (19). This may be even more significant in the case of *Drosophila* AGO2 that has a faster turnover rate than human AGO2 (20,21). Thus, for *Drosophila* AGO2 that can catalyze multiple rounds of RNA cleavage, availability of the target is certainly the limiting factor in silencing efficiency. The preferential association of AGO2 with ribosomes, already highlighted by others (17,18), suggests that this subcellular localization determines the preferential targeting of viral transcripts. We propose that the RISC complex is constantly present at translation sites, where it could most efficiently scan incoming viral mRNAs at the step where they are most exposed and before they are translated into viral proteins.

Our work has implications for strategies designed to target viral infection using dsRNA mediated silencing in insects and possibly other organisms. Optimal target design should consider genes that are more likely to affect viral replication. This may not be a major issue in the case of RNA viruses that often do not carry non-essential genes. Nevertheless, some genes may have more striking effects than others, especially the ones that are found in lower amounts in infected cells. In regard to positive stranded RNA viruses in which the genome may act directly as a transcript for translation, we did not test viruses that encode a single polyprotein, such as Flaviviruses, but this may be a case where genome and transcripts may be indistinguishable.

In conclusion, our results provide new insights on the control of viral infection by siRNAs in animals, which integrates well with the strategic struggle between the virus and the host cell for the control of ribosomes.

## Materials and Methods

### Virus stocks and cells

VSV-WT was a gift from Dr. Erna Geessien Kroon (strain Indiana isolate P94). VSV-GFP and SINV-GFP were a gift from Curt Horvath and Ilya Frolov, respectively. For viruses propagation, VERO cells were maintained on DMEM medium (LGC) supplemented with 10% heat-inactivated Fetal Bovine Serum (FBS, Gibco), 1XGlutaMAX (Gibco) and 1X penicillin/streptomycin (10mg.mL 1/10,000U,Gibco). Cells were seed to 80% of confluence and infected with VSV-WT or VSV-GFP at multiplicity of infection of 0.001 and for SINV-GFP, at 0.1. Cells were maintained for 20-36h at 37ºC until the appearance of cytopathic effects. After this time, the supernatant was collected and clarified by centrifugation to generate virus stocks that were kept at −80 °C before use. Mock-infected supernatants were prepared under the same condition without virus infection as control of experiments. Titration by plaque assay of VSV-WT and VSV-GFP was performed on VERO cells and for SINV-GFP on BHK-21 cells. VSV-WT inactivation was done by UV radiation at the given 254nm wavelength as previously described (22). VSV-UV was titrated after UV exposure to verify viral inactivation. *Drosophila* S2 cells were maintained in Schneider’s medium supplemented with 10% (FBS, Gibco), 1XGlutaMAX (Gibco), and 1X penicillin/streptomycin (10mg.mL 1/10,000U,Gibco). Cells were incubated at 28ºC for viral infections.

### *Drosophila* and S2 infections

Flies stocks were raised on standard cornmeal-agar medium at 25 °C. Adult females flies 3 to 6 days of age were used in infection experiments *in vivo. Canton S* genotype flies were obtained from the Bloomington Fly Stock Center (Bloomington, IN). Infections were done by intrathoracic injection (Nanoject II apparatus; Drummond Scientific) of a 69nL of 150ng of dsRNAs or viral suspension (5000 PFU/fly for VSV-WT and 500PFU/fly for VSV-GFP and SINV-GFP). Injected flies with dsRNAs or viruses were maintained at 25ºC for different times according to each experiment setup. After these times, flies were harvested in TRIzol reagent (Invitrogen). Samples were homogenized using a mini-Bead beater homogenizer (BioSpec). For *in vitro* experiments, 10^7^ S2 cells were counted, transferred to canonical tubes and infected with VSV-GFP at MOI of 1, at 28ºC under gentle homogenization. After 1h, cells were spun down at 200g, at 4ºC for 5min and the medium removed. Cells were resuspended in supplemented Schneider’s medium and seeded into 25-cm tissue culture dishes. Infected cells were incubated at 28ºC for 24h and submitted to cell fractionation.

### Quantification of viral RNAs by RT-qPCR

Total RNA from adult flies or S2 cells were extracted using Trizol reagent according to the manufacturer’s protocol (Invitrogen). 500ng of total RNA was reverse transcribed using 300 ng of random primers or 500 ng of anchored oligo dT primers. 2uL of diluted cDNA was used as template for qPCR reaction containing Sybr Green (Invitrogen) and specific primers. RT-qPCR was performed using QuantStudio 7 Flex™ RealTime PCR System (Applied® Biosystems). Analyses of RNA expression was done using ΔC_T_ method with Rpl32 as the internal control gene. Primer sequences are provided in **S1 Table**.

### dsRNA constructions and treatments

*In vitro* transcription was done using T7/SP6 MEGAscript Kits (Ambion), following the manufacturer’s instructions. Briefly, dsRNAs targeting viral sequences were produced from a T7 promoter sequences obtained by PCR amplification from TOPO plasmid (Thermo #K4575J10) containing purified PCR product corresponding to N region of VSV gene (region: 427 – 962) or to nsP4 region of SINV gene (region: 5920 – 6424). dsFluc and dsGFP were produced from a T7 and SP6 promoters sequences containing the firefly luciferase sequence from pGL3-Basic plasmid (Promega) and GFP PCR amplification from plasmid pDSAG (Addgene #62289) respectively. Adult female flies were intrathoracically injected with 69nL of a dsRNA solution (150ng/fly) diluted in annealing buffer (20mM Tris-HCl pH7.5, 100mM NaCl). dsRNAs were tracked *in vivo* for their efficiency **(S1 Fig)**. Primers sequences for dsRNAs construction are provided in **S1 Table**.

### Cell fractionation

Cell fractionation was performed as described by Hammond et al (18). Briefing, log-phase S2 cells were plated on 25-cm tissue culture dishes and were VSV or mock-infected. After 24h, cells were harvested in PBS containing 5mM EGTA and washed twice in cold PBS and once in hypotonic buffer (10 mM HEPES pH 7.3, 6 mM b-mercaptoethanol). Cells were suspended in 1mL of hypotonic buffer containing complete protease inhibitors (Protease Inhibitor Cocktail Tablets, EDTA-Free) and 0.5 units mL−1 of RNAseOUT (Invitrogen) and then disrupted in a douncer homogenizer with a type B pestle. The extract was centrifuged at 800g for 5min at 4ºC for cell clarification and the supernatant was subsequently centrifuged at 100,000g for 3h. The resulting pellet, containing ribosomes, was extracted in hypotonic buffer containing 1 mM MgCl2 and 400 mM KOAc. The cytosolic fraction (S100) and ribosome fraction (P100) were used to quantify proteins.

### Western blotting

After cell fractionation, S100, P100 and input were resuspended in 6X Laemmli SDS PAGE sample loading buffer. Total cellular protein for each sample was subjected to SDS-PAGE, followed by electroblotting onto nitrocellulose membranes. Membranes were blocked with 5% w/v BSA or 10% w/v nonfat dry milk in TBST buffer (150 mM NaCl, 10 mM Tris–HCl, pH 7.4, and 0.1% Tween 20) for 1 h and then incubated with anti-rabbit Rps15 (Abcam#168361), anti-rabbit AGO2 (Abcam#32381) or anti-guinea-pig VSV (raised against the recombinant protein comprising the 188 N-terminal amino acids of VSV RdRp by Protéogenix) antibodies in TBST buffer containing 3% w/v BSA or 5% w/v nonfat dry milk overnight at 4°C. Membranes were rinsed three times with TBST buffer and incubated with the related secondary peroxidase conjugated anti-IgG antibody diluted 1:5000 in TBST buffer containing 3% w/v BSA or 5% w/v nonfat dry milk for 1 h. Membranes were rinsed three times with wash buffer, incubated with ECL prime western blotting detection reagents, and scanned and analyzed by ImageQuant LAS 4000 (GE Healthcare).

### Data Analysis

GraphPad Prism™ software was used to analyze data for statistical significance in viral RNA loads compered to control groups. A two-tailed student t test was used to statistically analyze the data determined by unpaired *t*-test followed by Mann-Whitney post-test.

## Acknowledgements

The authors would like to thank members of Marques and Imler groups for discussion and suggestions.

## Supporting information

**S1 Text. Efficiency test of gene silencing by dsRNAs**.

**S1 table. Primers sequences utilized for RT-qPCR assays and dsRNAs constructions**.

**Fig S1. Efficiency test of gene silencing by dsRNAs. (A)** Experimental design of dsGFP treatment in *Hml-delta-GAL4, UAS-GFP* flies. **(B)** GFP RNA levels at day 3 post treatment with dsGFP. **(C)** Experimental design of dsVSV-N or dsSINV-nsp4 treatment and VSV- or SINV-GFP infections in *Canton S* flies. **(D)** VSV-L and SINV-nsp4 RNA levels at day 4 post infection with VSV- or SINV-GFP infection. mRNA levels were assessed by RT-qPCR performed in triplicate and normalized to Rpl32 mRNA levels. Violin plots show the frequency of data distribution with the median and quartiles. Each dot represents an individual sample. Numbers of samples are indicated above each violin plot. The dashed line represents the detection threshold of RT-qPCR. *p* values are shown for statistical significant *(p<0*.*005)* between dsFluc and dsGFP **(B)** and dsFluc comparing to dsVSV-N or dsSINV-nsp4 samples **(D)** determined by two-tailed Mann–Whitney U-test.

## References

1. Soares ZG, Gonçalves ANA, de Oliveira KPV, Marques JT. Viral RNA recognition by the Drosophila small interfering RNA pathway. Microbes Infect. 2014 Dec;16(12):1013–21.

2. Ding SW. RNA-based antiviral immunity. Nat Rev Immunol. 2010 Sep;10(9):632–44.

3. Li H, Li WX, Ding SW. Induction and suppression of RNA silencing by an animal virus. Science. 2002 May 17;296(5571):1319–21.

4. Galiana-Arnoux D, Dostert C, Schneemann A, Hoffmann JA, Imler JL. Essential function in vivo for Dicer-2 in host defense against RNA viruses in drosophila. Nat Immunol. 2006 Jun;7(6):590–7.

5. van Rij RP, Saleh MC, Berry B, Foo C, Houk A, Antoniewski C, et al. The RNA silencing endonuclease Argonaute 2 mediates specific antiviral immunity in Drosophila melanogaster. Genes Dev. 2006 Nov 1;20(21):2985–95.

6. Wang XH, Aliyari R, Li WX, Li HW, Kim K, Carthew R, et al. RNA interference directs innate immunity against viruses in adult Drosophila. Science. 2006 Apr 21;312(5772):452–4.

7. Marques JT, Wang JP, Wang X, Oliveira KPV de, Gao C, Aguiar ERGR, et al. Functional Specialization of the Small Interfering RNA Pathway in Response to Virus Infection. PLOS Pathog. 2013 Aug 29;9(8):e1003579.

8. Han YH, Luo YJ, Wu Q, Jovel J, Wang XH, Aliyari R, et al. RNA-Based Immunity Terminates Viral Infection in Adult Drosophila in the Absence of Viral Suppression of RNA Interference: Characterization of Viral Small Interfering RNA Populations in Wild-Type and Mutant Flies▿. J Virol. 2011 Dec;85(24):13153–63.

9. Mueller S, Gausson V, Vodovar N, Deddouche S, Troxler L, Perot J, et al. RNAi-mediated immunity provides strong protection against the negative-strand RNA vesicular stomatitis virus in Drosophila. Proc Natl Acad Sci U S A. 2010 Nov 9;107(45):19390–5.

10. Hong H, Wang C, Huang Y, Xu M, Yan J, Feng M, et al. Antiviral RISC mainly targets viral mRNA but not genomic RNA of tospovirus. PLoS Pathog. 2021 Jul;17(7):e1009757.

11. Banerjee AK. Transcription and replication of rhabdoviruses. Microbiol Rev. 1987 Mar;51(1):66–87.

12. Lim K il, Lang T, Lam V, Yin J. Model-Based Design of Growth-Attenuated Viruses. PLoS Comput Biol. 2006 Sep;2(9):e116.

13. Morin B, Rahmeh AA, Whelan SPJ. Mechanism of RNA synthesis initiation by the vesicular stomatitis virus polymerase. EMBO J. 2012 Mar 7;31(5):1320–9.

14. Barton DJ, Sawicki SG, Sawicki DL. Solubilization and immunoprecipitation of alphavirus replication complexes. J Virol. 1991 Mar;65(3):1496–506.

15. Rupp JC, Sokoloski KJ, Gebhart NN, Hardy RW. Alphavirus RNA synthesis and non-structural protein functions. J Gen Virol. 2015 Sep;96(9):2483–500.

16. Ishizuka A, Siomi MC, Siomi H. A Drosophila fragile X protein interacts with components of RNAi and ribosomal proteins. Genes Dev. 2002 Oct 1;16(19):2497–508.

17. Caudy AA, Myers M, Hannon GJ, Hammond SM. Fragile X-related protein and VIG associate with the RNA interference machinery. Genes Dev. 2002 Oct 1;16(19):2491–6.

18. Hammond SM, Bernstein E, Beach D, Hannon GJ. An RNA-directed nuclease mediates post-transcriptional gene silencing in Drosophila cells. Nature. 2000 Mar 16;404(6775):293–6.

19. Ruijtenberg S, Sonneveld S, Cui TJ, Logister I, de Steenwinkel D, Xiao Y, et al. mRNA structural dynamics shape Argonaute-target interactions. Nat Struct Mol Biol. 2020 Sep;27(9):790–801.

20. Haley B, Zamore PD. Kinetic analysis of the RNAi enzyme complex. Nat Struct Mol Biol. 2004 Jul;11(7):599–606.

21. Hutvágner G, Zamore PD. A microRNA in a multiple-turnover RNAi enzyme complex. Science. 2002 Sep 20;297(5589):2056–60.

22. Lytle CD, Sagripanti JL. Predicted inactivation of viruses of relevance to biodefense by solar radiation. J Virol. 2005 Nov;79(22):14244–52.

